# Glutamine supports human cytomegalovirus induced lipidome remodeling through reductive carboxylation

**DOI:** 10.64898/2026.01.26.701865

**Authors:** Rebekah L. Mokry, Caleb B. de Lacy, John G. Purdy

**Affiliations:** BIO5 Institute, University of Arizona, Tucson, Arizona, USA; Department of Immunobiology, University of Arizona, Tucson, Arizona, USA; Cancer Biology Interdisciplinary Program, University of Arizona, Tucson, Arizona, USA

## Abstract

Lipidome remodeling during human cytomegalovirus (HCMV) replication is a complex process that requires induction of lipogenic proteins and altered metabolite flow to support synthesis of fatty acids and lipids. HCMV infection increases the utilization of glucose and acetate to provide enough carbons to support increased demand for lipogenesis during virus replication, but other carbon contributors have not been studied. Here, we identify glutamine as a carbon source for lipogenesis during HCMV infection. Metabolic tracing with ^13^C-labeled glutamine revealed carbons from glutamine are enriched in phospholipids and neutral lipids during infection, including phosphatidylcholine, phosphatidylethanolamine, diacylglycerol, and triacylglycerol. Additional metabolic tracing demonstrates that HCMV infection promotes glutamine flow to fatty acid synthesis primarily through reductive carboxylation, i.e., conversion of glutamine to citrate through isocitrate. Through the use of two different ^13^C-labeled forms of glutamine, we found that ∼30% of the carbons from glutamine are delivered to fatty acid synthesis through additional metabolic means. Our current understanding of metabolite utilization during HCMV replication is based on cell culture models where there is an excess amount of glucose, suggesting that deriving carbons from glutamine might be needed when glucose levels are low. To determine if concentrations of glucose and glutamine change their contributions to fatty acid synthesis, we investigated lipogenesis when glucose and glutamine are at physiological levels (5 mM and 0.55 mM, respectively). We determined that physiological levels of glucose and glutamine are sufficient to support the increased demand for fatty acid synthesis caused by HCMV infection, despite a reduction in virus production. Using metabolic tracing with ^13^C-labeled forms of glucose or glutamine, we determined that both carbon sources still contribute to fatty acid synthesis when present at physiological levels. Overall, our results identify viral activation of reductive carboxylation that increases glutamine flow to lipogenesis during infection. This work provides additional insight into metabolic reprogramming that supports HCMV-induced lipidome remodeling.

**Author Summary:** Many viruses hijack cellular metabolic processing to obtain the components needed for replication. Human cytomegalovirus (HCMV) uses several mechanisms to reprogram lipid metabolism and remodel the lipidome of infected cells. HCMV promotes synthesis of very long chain fatty acids that are found in phospholipids and triacylglycerol. Glucose and acetate contribute carbon to fatty acid synthesis and elongation following HCMV infection. In this work, we demonstrate that glutamine is an additional carbon source for fatty acid and lipid synthesis. Phospholipids and neutral lipids are enriched with carbons from glutamine during HCMV infection. Mechanistically, HCMV induces reductive carboxylation to increase glutamine flow to fatty acid synthesis and increased metabolite availability supports additional carbon flow to fatty acids. Overall, this study provides additional insight into virus-induced metabolic remodeling that supplies the molecular building blocks for virus replication.

## Introduction

Viruses depend on the metabolic environment of their host cell to successfully replicate. Many viruses manipulate metabolic pathways to drive synthesis of molecular building blocks for replication such as DNA, protein, and lipids. Nutrients in the environment play specific roles in viral reprogramming of metabolism during infection. Two major nutrients, glucose and glutamine, are crucial for replication of several viruses, including the beta-herpesvirus human cytomegalovirus (HCMV) (1). HCMV is a prevalent virus that is the leading cause of viral-mediated congenital disorders, including microcephaly, developmental and cognitive disabilities, and hearing loss (2, 3). In immunocompromised individuals, such as those who have undergone solid-organ or stem cell transplant or those with secondary immune suppressive infections, HCMV can cause severe, life-threatening diseases (4). HCMV has evolved an adept system to manipulate and reprogram cellular metabolism and provides a model to understand the mechanisms by which viruses hijack and rewire nutrient flow (1, 5).

HCMV requires lipid synthesis for efficient replication and remodels the lipidome during infection (6–11). Lipids support viral assembly and infectivity of the viral particle (11–13). HCMV remodels the lipidome in part by promoting fatty acid (FA) synthesis and elongation to increase very long chain FA (VLCFA) with 26 or more carbons (10, 11). VLCFA are used to generate phospholipids and neutral lipids with very long chain fatty acyl tails (7–9). To promote lipid synthesis, HCMV induces lipogenic proteins involved in synthesis and elongation of FA and drives glucose carbons to citrate and acetyl-CoA to support FA synthesis (6, 7, 10, 11, 14, 15). Since glucose carbons are fueling FA production, glutamine uptake and glutaminolysis, i.e., catabolism of glutamine to support energy production and synthesis of other macromolecules, are induced to support the TCA cycle (16).

Glutamine has been largely overlooked as a potential carbon contributor for FA and lipid synthesis since glucose is the primary source of carbon for FA synthesis (6, 10, 11). It is well established that glutamine maintains the TCA cycle during HCMV infection (6, 16), but other roles of glutamine during infection are not well defined (17). In this study, we identify glutamine as a carbon source for lipogenesis during HCMV replication. Using metabolic tracing coupled with liquid chromatography high-resolution tandem mass spectrometry (LC-MS/MS), we found that glutamine directly contributes to lipid synthesis. We determined that glutamine supports lipogenesis of most lipid classes included in this study. Additionally, glutamine supports synthesis of lipids across lipid tail lengths (number of carbons) and degrees of desaturations (number of double bonds). Moreover, we found that HCMV infection induces reductive carboxylation, i.e., “reverse” metabolism of glutamine to citrate, during which glutamine is metabolized to α-ketoglutarate, converted to isocitrate, and then to citrate, which supplies acetyl-CoA for FA synthesis. Although reductive carboxylation is the primary mechanism through which glutamine supports FA synthesis, we identified additional glutamine carbon flow to FA production that is likely supported by malic enzyme pathway. Finally, to understand the potential effect to HCMV lipogenesis of glucose and glutamine availability in the nutrient environment, we reduced the high level of glucose and glutamine found in standard DMEM to physiological concentrations. When glucose and glutamine are provided at the level found in human plasma of healthy people, HCMV was able to promote FA synthesis and elongation but had a reduced amount of virus production. Overall, our data reveal that reductive carboxylation supports lipidome remodeling during HCMV replication even when culturing in physiological levels of glucose and glutamine. The findings expand our understanding of the role of glutamine during viral infection and demonstrate another example of HCMV-induced metabolic reprogramming that supports successful virus replication.

## Results

### Glutamine supports lipid and fatty acid synthesis during HCMV replication

Glutamine is required for HCMV replication (16, 17). HCMV increases glutaminolysis to support TCA cycle continuation (6, 16). Since glutamine can feed lipid synthesis outside the context of virus infection (18–21), we hypothesized that glutamine could support lipid synthesis during HCMV infection. To determine if glutamine supports lipogenesis, we used an isotopic labeling approach with uniformly labeled ^13^C glutamine (U-^13^C-Gln). Experiments were performed on fully confluent human foreskin fibroblast (HFFs) cultured in serum-free DMEM to eliminate any contribution of unlabeled glutamine in the bovine serum. These conditions are widely used in studies of HCMV metabolism (6–8, 10, 11, 14, 22–29). We infected or mock-infected HFFs at MOI 3. At 1 hpi, inoculum was removed, cells were washed with PBS, and DMEM containing 4 mM U-^13^C-Gln was provided to cells. At 48 and 72 hpi, lipids were extracted and identified using LC-MS/MS. These times were selected because HCMV induces lipidome remodeling by 48 hpi with more pronounced alterations by 72 hpi (7–9). We first examined if HCMV infection promotes glutamine flow to synthesis of the major lipid classes. To do so, the percentage of lipids containing at least one ^13^C atom were quantified after correcting for natural isotopic abundance. Each class of lipids has several dozen lipid species. We calculated the percent of labeled lipids by averaging the amount of label in the observed species within a lipid class. We also examined lysoPC (LPC) and lysoPE (LPE) that contain one fatty acyl tail since interconversion between one-tailed LPC or LPE with two-tailed PC or PE is active in fibroblasts (9, 30). By 48 hpi, the percentage of PC, PE, PI, DG, and TG labeled carbons were increased in HCMV-infected cells compared to mock (**Fig 1A**) and by 72 hpi these lipids had more pronounced labeling (**Fig 1B**). At 72 hpi, glutamine carbon labeling of PS, LPC, and LPE were also significantly increased during infection (**Fig 1B**). Most lipids from HCMV-infected cells had approximately two times more labeling from glutamine compared to mock by 72 hpi. We observed 8% in mock compared to 23% in HCMV-infected cells for PC, 6% compared to 16% for PE, 12% compared to 19% for PI, and 12% compared to 17% for PS. Labeled DG and TGs were a respective 11 and 10% in mock compared to 25 and 23% in infected cells. Labeled LPC and LPE were subtly, but significantly, increased during HCMV infection. LPC had 7% labeling in mock compared to 10% during infection and LPE had 1% labeled in mock compared to 2% during infection. PG was the only lipid class that we examined where infection did not alter glutamine labeling (**Fig 1A-B**).

**Fig 1.**
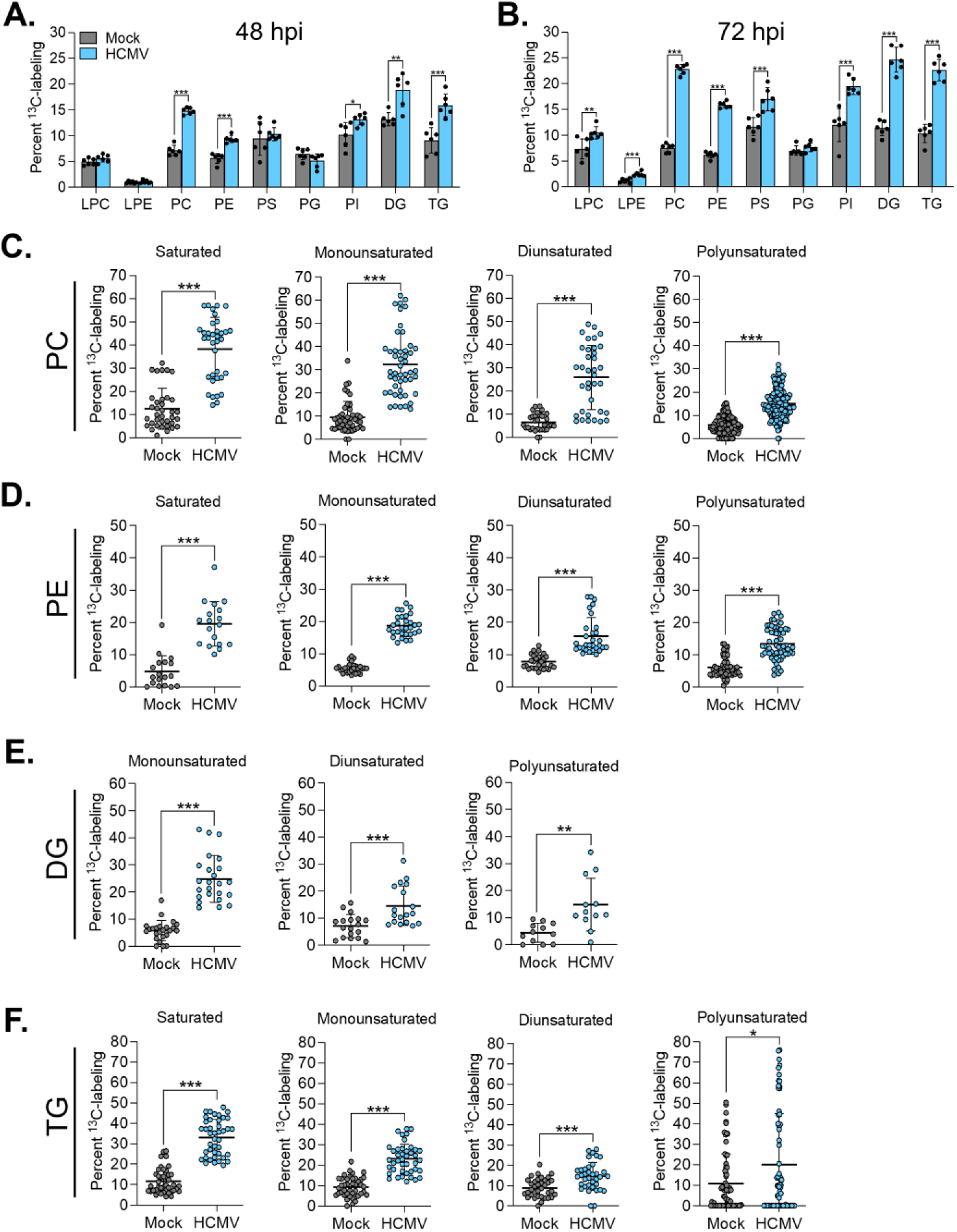
Glutamine supports lipid synthesis during HCMV replication. Human foreskin fibroblasts (HFFs) were infected with TB40/E-GFP or mock-infected. At 1 hpi, virus inoculum was removed and replaced with media containing uniformly labeled ^13^C glutamine (U-^13^C-Gln). At 48 and 72 hpi, lipids were extracted and measured using LC-MS/MS. (**A-B**) Percent of ^13^C-labeled lipids were calculated for each lipid class by averaging the percent label for each lipid species at 48 (**A**) and 72 hpi (**B**). Percent of ^13^C-labeled lipids by tail desaturations for PC (**C**), PE (**E**), DG (**E**), and TG (**F**) at 72 hpi. Each dot represents a lipid from one replicate. Error bars represent standard deviation (SD). Unpaired t-test were used to determine significance. *P* < 0.05, *; *P* < 0.01, **; *P* < 0.001, ***. *n*=6

Glutamine carbon contribution to lipid synthesis could be directed to fatty acyl tails of the lipids. To determine if glutamine supports synthesis of different tail types during infection, we quantified the percent of labeled lipids containing saturated (no double bond), monounsaturated (one double bond), diunsaturated (two double bonds), or polyunsaturated (three or more double bonds) fatty acyl tails at 72 hpi (**Fig 1C-F**). For lipids that have two fatty acyl tails, saturated contain two saturated tails, while monounsaturated lipids contain one saturated and one monounsaturated tail. Diunsaturated lipids contain one saturated tail and one diunsaturated tail or two monounsaturated tails, and polyunsaturated lipids contain one saturated or monounsaturated tail with a polyunsaturated tail or two polyunsaturated tails. We focused analysis on lipid classes that showed the highest levels of labeling from glutamine in HCMV-infected cells compared to mock, i.e., PC, PE, and DG. For labeled PC and PE, HCMV significantly increased the percent ^13^C in all tail types with greatest enrichment for saturated, monounsaturated, and diunsaturated (**Figs 1C-D** and **S1A-B**). Labeled DG tails were also increased during HCMV infection with the greatest enrichment for monounsaturated tails (**Figs 1E** and **S1C**). Saturated DG were not identified in this experiment. We also quantified ^13^C-labeling of TG tail types which contain three fatty acyl tails. Monounsaturated TG contain two saturated and one monounsaturated tail, diunsaturated could contain one saturated and two monounsaturated tails or two saturated and one diunsaturated tail. We observed the highest labeling for saturated TG tails during infection with mono- and diunsaturated also increased compared to mock (**Figs 1F** and **S1D**). Polyunsaturated TG could contain several different tail combinations, including three monounsaturated tails or two unsaturated and one polyunsaturated tail. We observed wide variability in the percentage of labeling for polyunsaturated TG, which may be due to the broad range of tail combinations that comprise TG lipids (**Figs 1F** and **S1D**). Overall, these data demonstrate that HCMV infection promotes glutamine use for lipid synthesis during infection.

The results of labeling lipids using U-^13^C-Gln suggest that glutamine is utilized for FA synthesis during infection (**Fig 1**). To directly test if glutamine is a carbon source for FA synthesis, we measured FA containing carbons from glutamine. In these experiments, esterified FA were liberated from lipids via saponification before measurement by LC-MS. This method enables us to measure all esterified tails independent of lipid class to gain a better understanding of how glutamine contributes to the building blocks used in lipid synthesis. We infected or mock-infected HFFs and cultured in media containing U-^13^C-Gln. FA were extracted at 48 and 72 hpi and identified using LC-MS. FA with at least one ^13^C were quantified after correcting for natural isotopic abundance. Since glutamine primarily contributes to FA synthesis through its conversion to citrate which then yields acetyl-CoA, the U-^13^C-Gln labeling strategy is expected to yield two ^13^C atoms per two-carbon unit used for FA synthesis and elongation (**Fig 2A**). We focused our analysis on saturated (SFA) and monounsaturated FA (MUFA) since HCMV strongly induces their production (10, 11), and our lipid labeling indicate that lipids with saturated and monounsaturated fatty acyl tails have the highest percentage of label (**Fig 1C-F**). In mock-infected cells, low levels of labeled SFA and MUFA were observed at both 48 and 72 hpi (**Figs 2B-C** and **S2A-B**). In HCMV-infected cells, labeled FA were increased by 48 hpi compared to mock and labeling were further enhanced by 72 hpi. By 72 hpi, labeled FA were significantly increased in HCMV-infected cells for all identified SFA and very long chain MUFA beginning with C24:1 (**Figs 2B-C**).

**Fig 2.**
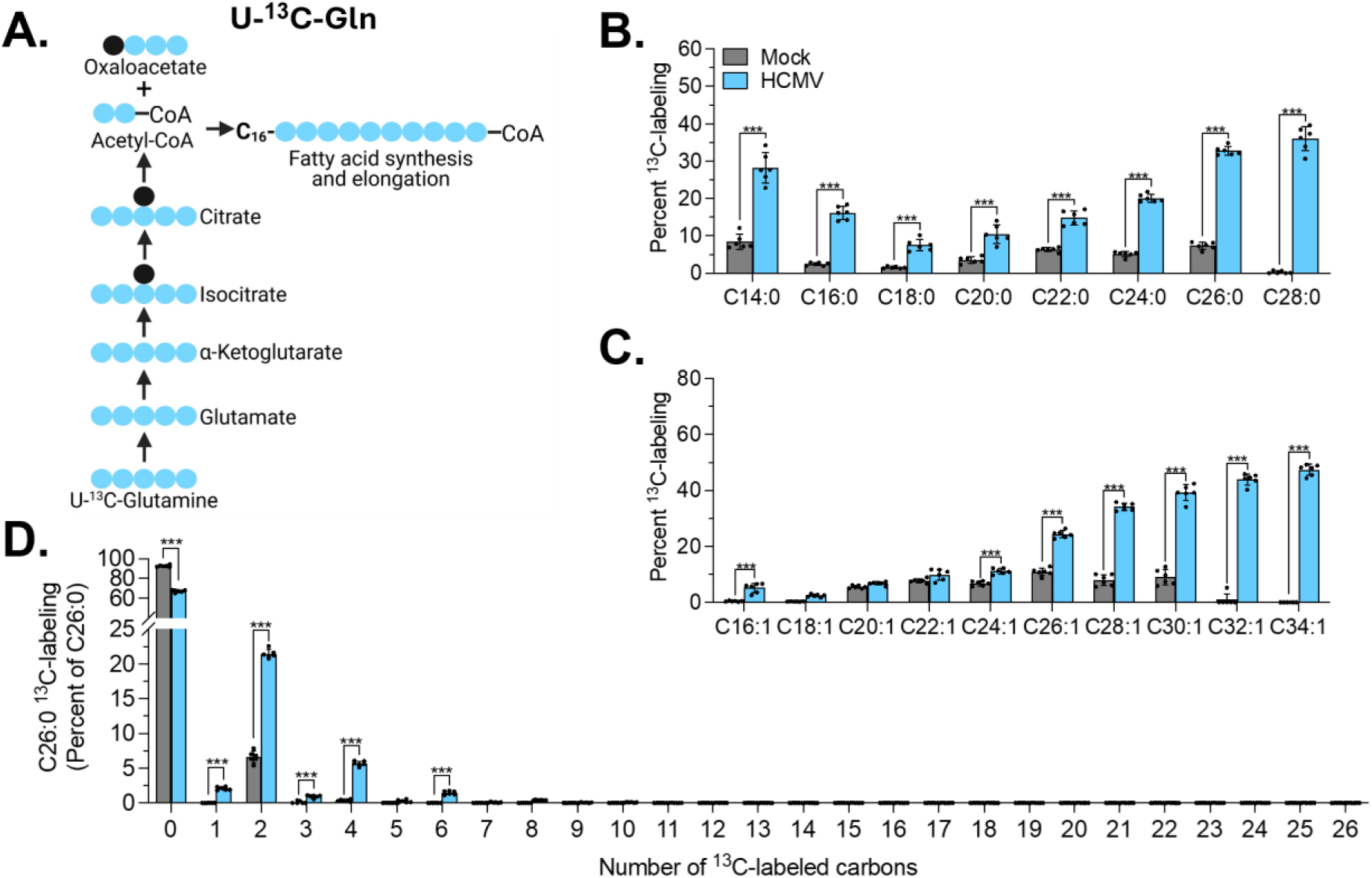
Glutamine supports fatty acid synthesis during HCMV replication. (**A**) Schematic of predicted fatty acid (FA) labeling from U-^13^C-Gln. Blue circles indicate ^13^C-labeled carbons, and black circles indicate unlabeled carbons. (**B-D**) HFFs were infected with TB40/E-GFP or mock-infected. At 1 hpi, virus inoculum was removed and replaced with media containing U-^13^C-Gln. At 72 hpi, lipids were extracted and FA were saponified. Labeled FA were measured using LC-MS. Percent ^13^C-labeled SFA (**B**) and MUFA (**C**) were quantified. (**D**) Percent ^13^C-labeled C26:0. Error bars represent SD. Two-way ANOVA with Šídák’s test were used to determine significance. *P* < 0.001, ***. *n*=6

Our analyses to this point focused on lipids or FA that contain one or more ^13^C atoms from glutamine. Next, we quantified the number of carbons incorporated into FA to better understand the extent of carbons in FA coming from glutamine. We focused on C26:0 since it showed a high level of labeling from glutamine as displayed in **Fig 2B** and has been previously examined for glucose and acetate labeling (7, 11). By 72 hpi, C26:0 in HCMV-infected cells have up to 8-labeled atoms derived from glutamine, whereas mock-infected cells contained only up to 4 labeled atoms (**Fig 2D**). Of note, we observed a small amount of 1-labeled C26:0, suggesting that labeled carbons may be entering FA synthesis through additional steps. Overall, our data demonstrate that glutamine contributes to FA synthesis and elongation during HCMV replication.

### Reductive carboxylation supports FA synthesis during HCMV replication

Glutamine primarily contributes to FA synthesis via reductive carboxylation, also called reductive glutamine metabolism, which metabolizes glutamine to citrate through the intermediate α-ketoglutarate in a series of enzymatic reactions (**Fig 2A**) (18–21). This pathway requires the activity of isocitrate dehydrogenase 1 (IDH1) or isocitrate dehydrogenase 2 (IDH2) that catalyze the conversion of α-ketoglutarate to isocitrate in a NADPH-dependent manner (**Fig 3A**). Both enzymes can also support the reverse reaction. This α-ketoglutarate to isocitrate reaction can occur in the cytosol (IDH1) or mitochondria (IDH2) (18–20, 31, 32). We investigated IDH1 and IDH2 levels to determine if these enzymes are increased to support glutamine flow to citrate during HCMV infection. Neither IDH1 nor IDH2 were increased during HCMV replication despite increased glutamine utilization for FA and lipid synthesis (**Fig 3B-D**). These data suggest increased enzyme levels are not required for induction of glutamine flow to lipids during HCMV replication.

**Fig 3.**
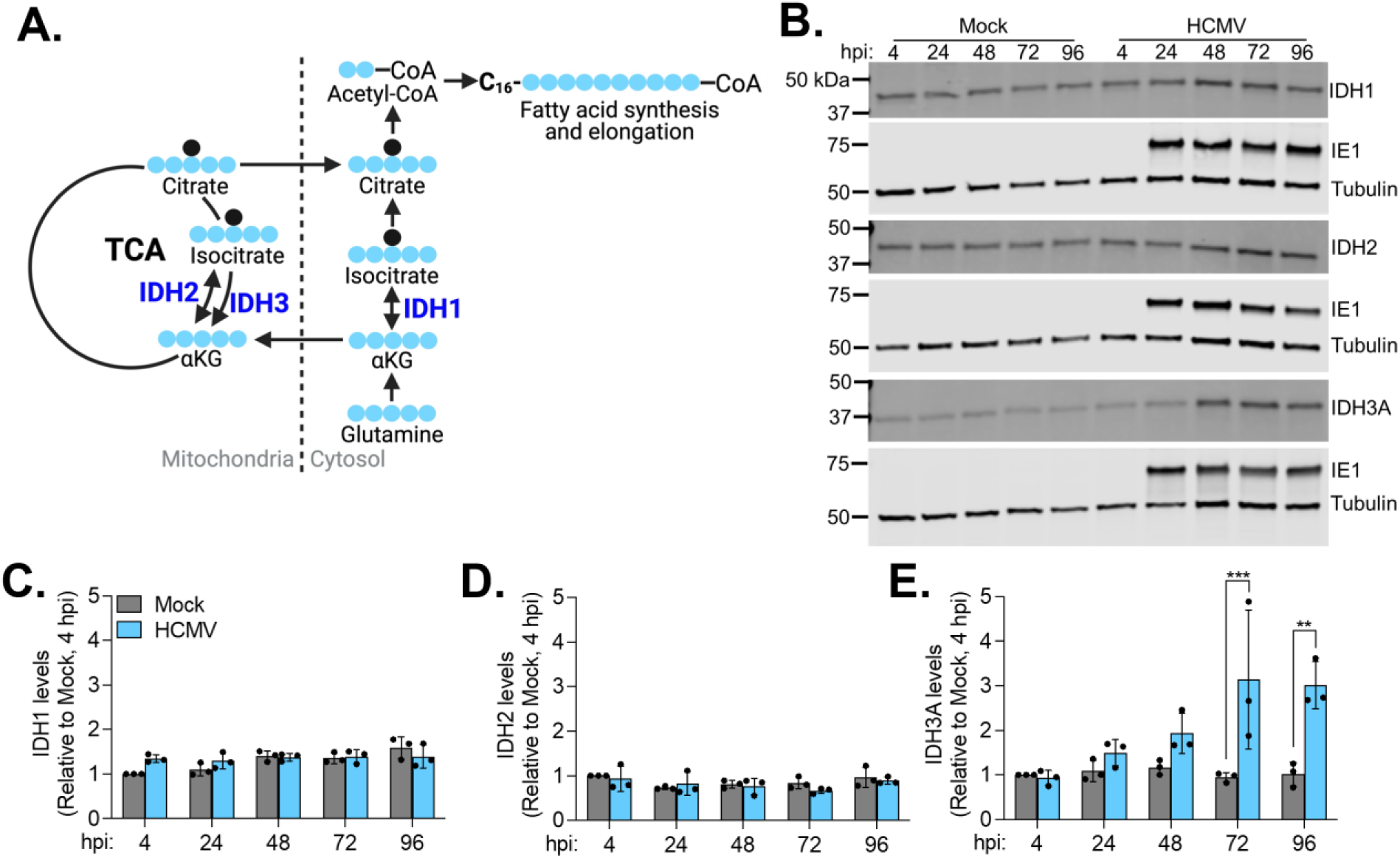
HCMV induces IDH3A but not IDH1 or IDH2. (**A**) Schematic of predicted FA labeling from U-^13^C-Gln with enzymes in bolded blue. Blue circles indicate ^13^C-labeled carbons, and black circles indicate unlabeled carbons. (**B-D**) HFFs were infected with TB40/E-GFP or mock-infected. Whole-cell lysates were collected at the indicated times points and analyzed by western blot. A representative western blot from three biological replicates is shown in **B**. Levels of IDH1 (**C**), IDH2 (**D**), or IDH3A (**E**) were normalized to tubulin and quantified relative to mock at 4 hpi. Error bars represent SD. Two-way ANOVA with Šídák’s test were used to determine significance. *P* < 0.01, **; *P* < 0.001, ***. *n*=3

Isocitrate dehydrogenase 3 (IDH3) supports oxidative turning of the TCA cycle and isocitrate conversion to α-ketoglutarate in an NAD^+^ dependent manner. We investigated IDH3 levels by blotting for the subunit IDH3A. IDH3A is increased by 72 hpi during HCMV replication (**Fig 3B** and **E**), suggesting IDH3 is induced to support TCA cycle maintenance (6, 27). These data indicate that glutamine contribution to lipid synthesis does not require increased IDH1 and IDH2 levels, while increased IDH3 supports HCMV-induced TCA cycle activity (6, 27).

Next, we determined if reductive carboxylation is the primary pathway by which glutamine contributes to FA synthesis during HCMV replication. We infected or mock-infected HFFs and cultured them in media containing glutamine with ^13^C at position 5 (5-^13^C-Gln). Metabolism of 5-^13^C-Gln retains the labeled carbon on one atom of acetyl-CoA, thereby incorporating one ^13^C atom per two-carbon unit added to the FA chain (**Fig 4A**). In mock-infected cells, low levels of labeling were observed for SFA and MUFA at 48 and 72 hpi (**Figs 4B-C** and **S3A-B**). At 48 hpi, HCMV-infected cells showed increased labeling for C14:0, C16:0, and VLCFA compared to mock (**S3A-B).** By 72 hpi, labeled FA were significantly increased for all identified SFA and very long chain MUFA beginning with C26:1 (**Fig 4B-C**). For the labeled carbons of C26:0, HCMV-infected cells showed increased levels for 1 to 3-labeled atoms compared to mock-infected cells at 72 hpi (**Fig 4D**). These data confirm our conclusion that HCMV infection promotes glutamine use for FA synthesis and demonstrate glutamine contributes through reductive carboxylation.

**Fig 4.**
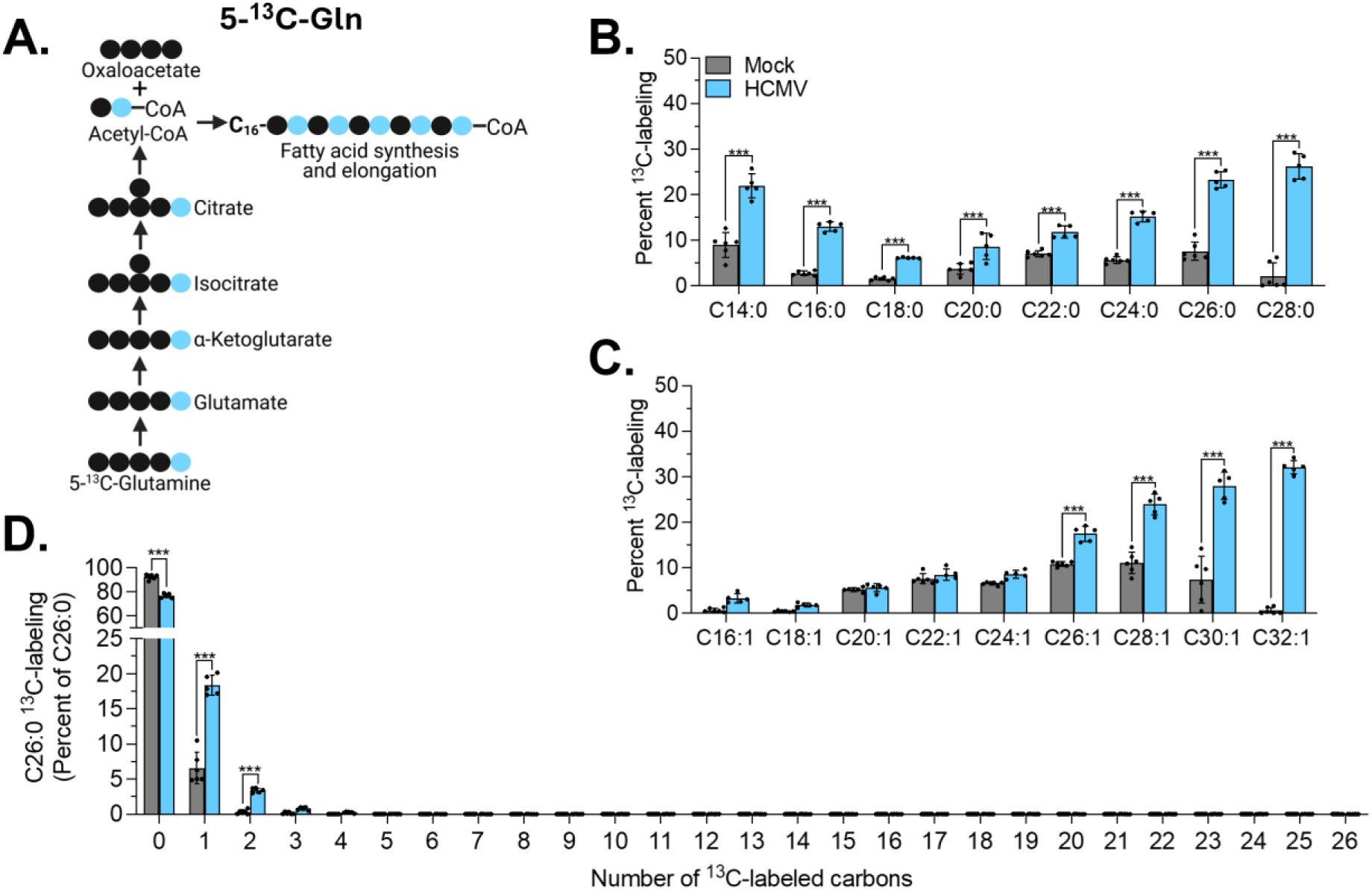
Glutamine is metabolized via reductive carboxylation during HCMV replication. (**A**) Schematic of predicted FA labeling from 5-^13^C-labeled glutamine (5-^13^C-Gln). Blue circles indicate ^13^C-labeled carbons, and black circles indicate unlabeled carbons. (**B-D**) HFFs were infected with TB40/E-GFP or mock-infected. At 1 hpi, virus inoculum was removed and replaced with media containing 5-^13^C-Gln. At 72 hpi, lipids were extracted and FA were saponified. Labeled FA were measured using LC-MS. Percent ^13^C-labeled SFA (**B**) and MUFA (**C**) were quantified. (**D**) Percent ^13^C-labeled C26:0. Error bars represent SD. Two-way ANOVA with Šídák’s test were used to determine significance. *P* < 0.001, ***. *n*=5-6

### Additional metabolic activity supports glutamine feeding FA synthesis during HCMV replication

Our glutamine labeling strategies provide information on the contribution of glutamine from reductive carboxylation (5-^13^C-Gln) versus the total contribution of glutamine (U-^13^C-Gln) to FA synthesis (18). Similar percentages of labeled FA would indicate that glutamine contributes to FA synthesis solely through reductive carboxylation. However, if the percentage of labeled FA is higher for U-^13^C-Gln labeling than 5-^13^C-Gln, it indicates glutamine contributes through additional pathways. To determine if additional pathways contribute to glutamine use for FA synthesis, we compared percentages of labeled SFA and MUFA from mock and HCMV-infected cultures from the two labeling strategies. Mock-infected cells showed similar percentage of labeled SFA and MUFA between the two labeled forms except for C28:1 at 72 hpi which showed a subtle increase in 5-^13^C-Gln fed cells (**Fig 5A-B**). These results indicate that the low level of FA labeling in mock-infected cells occurs through reductive carboxylation. In contrast, HCMV-infected cells showed significantly higher labeling from U-^13^C-Gln for SFA C14:0, C24:0, C26:0, and C28:0 (**Fig 5C**). Similar results were observed for MUFA with increased labeling of C26:1, C28:1, C30:1, and C32:1 (**Fig 5D**). These data indicate that reductive carboxylation accounts for the majority of FA labeling in HCMV-infected cells, but an additional pathway also contributes to glutamine flow to FA synthesis. To determine the percent of label from reductive carboxylation versus other pathways, we calculated the percent of FA label from 5-^13^C-Gln relative to U-^13^C-Gln. Labeling from U-^13^C-Gln was set to 100% and considered total FA labeling. For significantly altered SFA and MUFA between the two labeled glutamine forms, we found ∼70% of FA label occurs through reductive carboxylation (**Fig 5E-F**). These data demonstrate that glutamine supports FA synthesis primarily through reductive carboxylation during infection. However, ∼30% of FA label from glutamine is shuttled from additional metabolic steps during HCMV replication.

**Fig 5.**
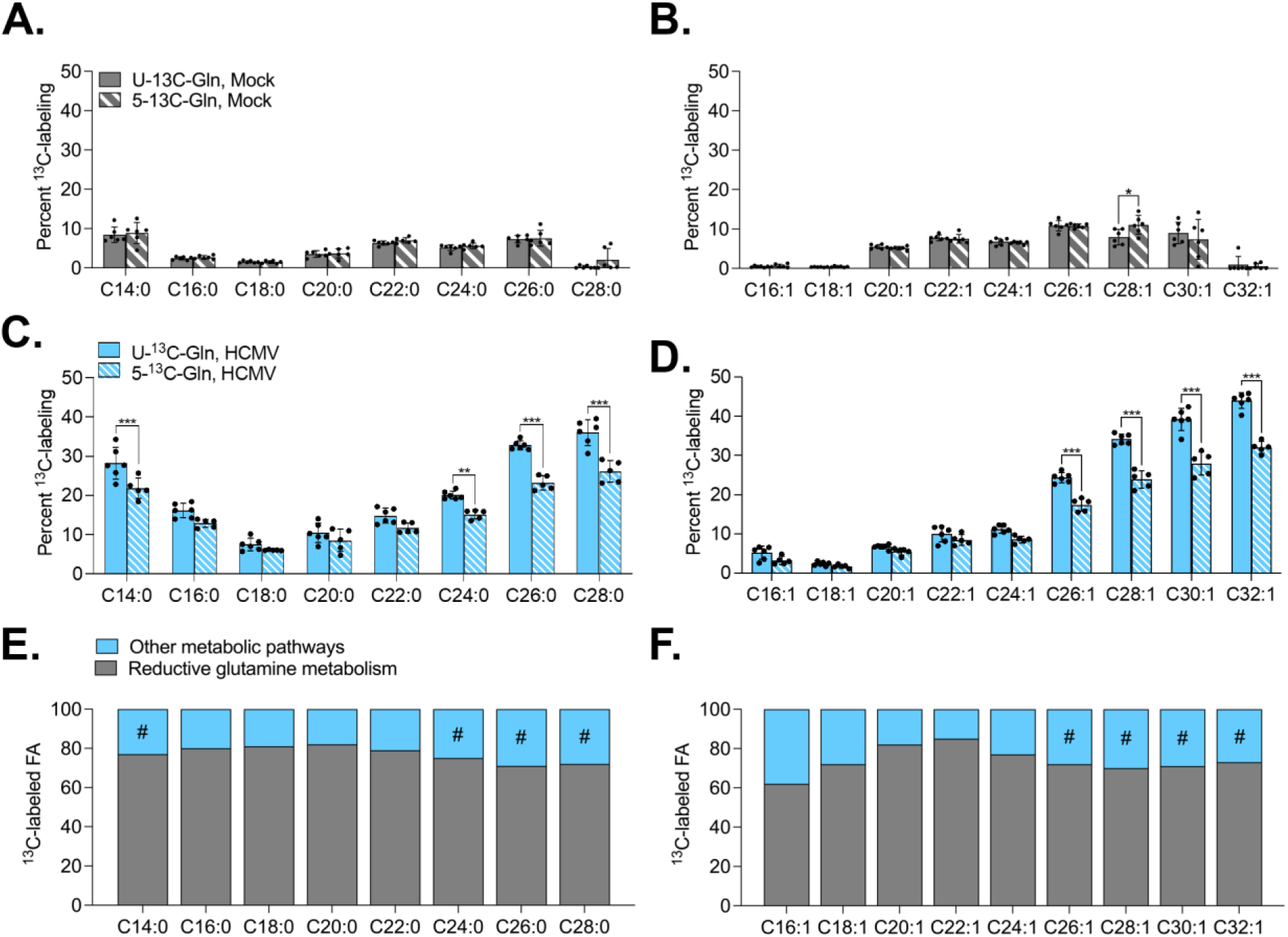
Reductive carboxylation is the primary pathway through which glutamine is metabolized to support FA synthesis. Percent ^13^C-labeled SFA (**A & C**) and MUFA (**B & D**) in mock or HCMV-infected cells at 72 hpi from **Fig 2B-C** and **4B-C** were compared. (**B & D**) Percent labeled FA in HCMV-infected cells from reductive carboxylation versus other pathways for SFA (**C**) and MUFA (**D**) were calculated by quantifying the ratio of 5-13-C to U-^13^C-Gln. Octothorpe (#) indicates statistically significant differences of U-^13^C-Gln compared to 5-^13^C-Gln shown in **C** and **D**. Error bars represent SD. Two-way ANOVA with Šídák’s test were used to determine significance. *P* < 0.01, **; *P* < 0.001, ***. *n*=5-6

Reductive carboxylation of glutamine yields cytosolic oxaloacetate when citrate is converted to acetyl-CoA via ATP Citrate Lyase (ACLY) (**Fig 6A**). Several fates are possible for oxaloacetate, including conversion to malate followed by conversion to pyruvate. Pyruvate can feed into the TCA cycle to generate additional citrate that can support FA synthesis. In the U-^13^C-Gln labeling approach, this metabolism would result in 2-labeled and 3-labeled forms of pyruvate that can feed oxidative TCA cycle. Each of these labeled forms will result in 2-labeled acetyl-CoA that can produce 2-labeled citrate via the TCA cycle. In this pathway, labeled citrate is exported to the cytosol and converted to unlabeled oxaloacetate and 2-labeled acetyl-CoA that can provide two ^13^C-labeled two-carbon units for FA synthesis (**Fig 6A**) (7). Additionally, glutamine is also feeding oxidative TCA cycle during HCMV infection (**Fig 6B**) (6, 16). U-^13^C-Gln feeding oxidative TCA would produce 4-labeled citrate when combined with unlabeled acetyl-CoA from glucose or 6-labeled citrate if combined with acetyl-CoA from the pathway described above. Citrate is exported to the cytosol and converted to 4-labeled oxaloacetate and acetyl-CoA. Similar to oxaloacetate derived from reductive carboxylation, conversion to malate and then pyruvate would yield 3-labeled pyruvate that can feed oxidative TCA and produce 2-labeled acetyl-CoA for FA synthesis as described previously (**Fig 6B**). Cytosolic oxaloacetate produced from oxidative TCA-derived and reductive carboxylation-derived citrate converted to malate and then pyruvate would account for the additional labeling observed in U-^13^C-Gln compared to 5-^13^C-Gln (**Figs 5C-F** and **6A-B**).

**Fig 6.**
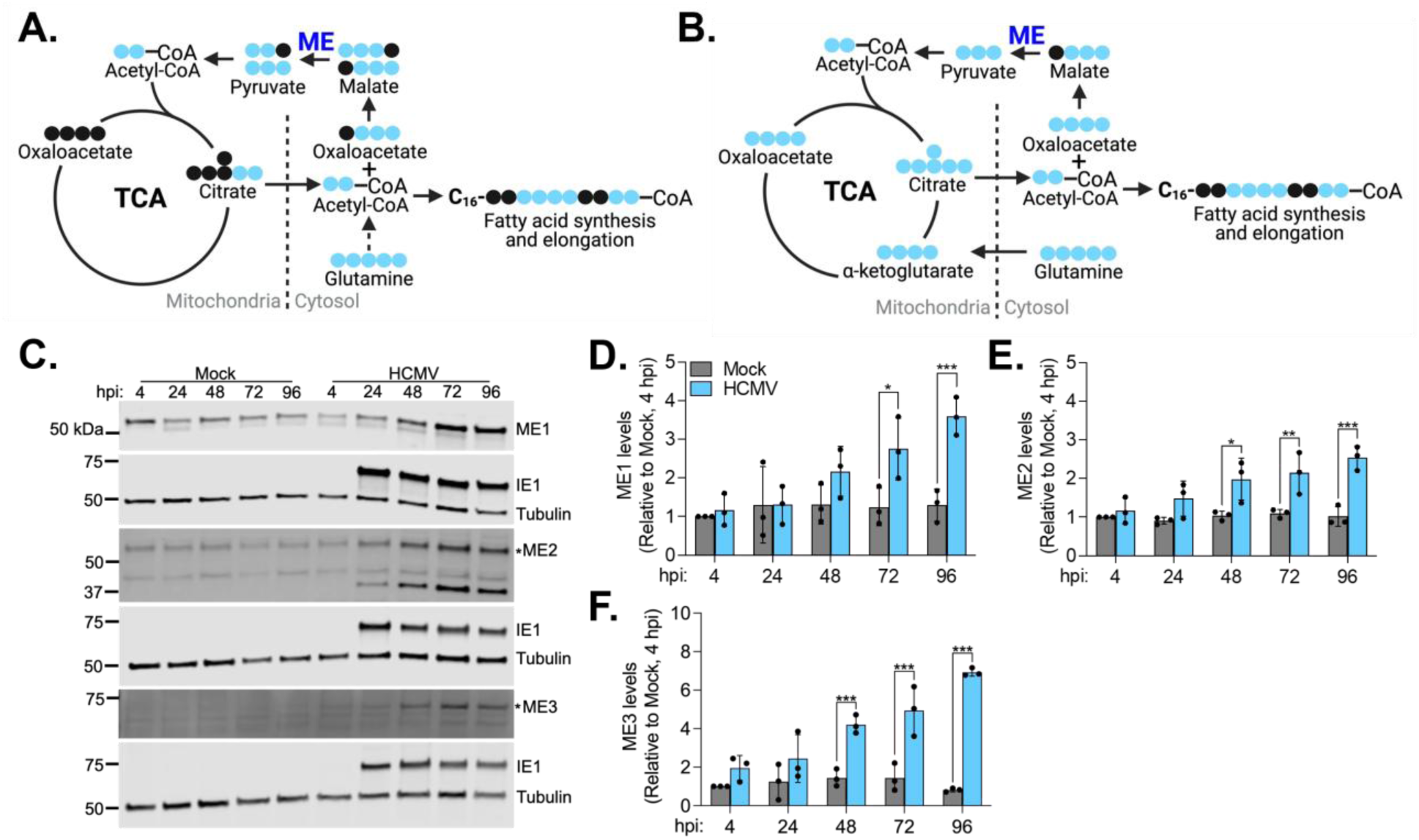
HCMV induces malic enzymes. (**A-B**) Schematic of predicted FA labeling from malic enzyme pathway using U-^13^C-Gln. Blue circles indicate ^13^C-labeled carbons, and black circles indicate unlabeled carbons. Enzymes are in bright blue. (**C-F**) HFFs were infected with TB40/E-GFP or mock-infected. Whole-cell lysates were collected at the indicated times and analyzed by western blot. A representative western blot from three biological replicates is shown in **B**. Levels of ME1 (**C**), ME2 (**D**), or ME3 (**E**) were normalized to tubulin and quantified relative to mock at 4 hpi. Asterisk (*) indicates protein band based on reported protein size. Error bars represent SD. Two-way ANOVA with Šídák’s test were used to determine significance. *P* < 0.01, **; *P* < 0.001, ***. *n*=3

Malate conversion to pyruvate is catalyzed by malic enzyme (ME). There are three ME isoforms: ME1 and ME3 are NADP^+^-dependent and found in the cytosol and the mitochondria, respectively, whereas ME2 is localized to the mitochondria with dual NAD^+^ and NADP^+^ specificity but a preference for NAD^+^ (33–37). We measured ME1, ME2, and ME3 to determine if HCMV infection alters their levels. ME1 levels are increased by 72 hpi, and ME2 and ME3 levels are increased by 48 hpi during HCMV replication (**Fig 6C-F**), suggesting HCMV induces these enzyme levels to support malate conversion to pyruvate. Collectively, increased ME levels coupled with U-^13^C-Gln versus 5-^13^C-Gln FA labeling results suggest that increased cytosolic oxaloacetate availability produced from citrate derived from oxidative TCA cycle and reductive carboxylation could feed the malic enzyme pathway and supply additional carbons to support HCMV-induced FA synthesis (**Figs 5C-F** and **6**).

### Physiological levels of glutamine and glucose support increased FA levels during HCMV replication

Our results demonstrate that glutamine feeds lipid and FA synthesis during HCMV replication, including through reductive carboxylation and the actions of ME. Thus far, our analysis of metabolite use for FA synthesis used nutrient concentrations found in DMEM as is common for studies investigating HCMV interactions with host metabolism (6, 11, 14, 16, 22, 24, 38, 39). The concentration of glucose and glutamine is at supraphysiological levels in most cell culture media, including DMEM. Next, we sought to determine if physiological levels of glutamine alter its overall contribution to FA synthesis. In these experiments, we also reduced the level of glucose to human sera concentrations, since glucose was found to be the primary carbon source for FA synthesis in HCMV infection (6, 7, 10, 11). First, we determined if infection altered the levels of FA in DMEM with physiological glucose (5 mM) and glutamine (0.55 mM) compared to standard DMEM (25 mM glucose and 4 mM glutamine). We chose these physiological concentrations from levels reported in healthy human sera and are used in cell culture media that mimic nutrient levels in human plasma (40, 41). We infected or mock-infected HFF-1 cells and changed the media at 1 hpi to standard DMEM with 25 mM glucose and 4 mM glutamine (25/4) or DMEM containing 5 mM glucose and 0.55 mM glutamine (5/0.55). FA were measured at 72 hpi. In mock-infected cells, reducing glucose and glutamine resulted in little change to FA levels with only small differences in VLCFA that are at low abundance in uninfected cells (**Fig 7A**). As expected, HCMV infection in standard 25/4 DMEM increased the levels of VLCFA relative to either mock condition. HCMV replication in 5/0.55 DMEM similarly increased VLCFA levels, indicating that physiological levels of glucose and glutamine are sufficient to support increased FA synthesis activity promoted by infection. Since our earlier experiments showed an increase in glutamine labeling of saturated and monounsaturated VLCFA, we examined the levels of C26:0, C26:1, C28:0, and C28:1 in cells cultured in 25/4 and 5/0.55. In mock-infected cells, these VLCFA levels in 5/0.55 remained similar to those cultured in 25/4 (**Fig 7B-E**). As anticipated, these four VLCFA were increased in HCMV-infected cells cultured in 25/4. Further, levels in infected cells cultured in 5/0.55 were similar in abundance to infected cells cultured in high glucose and glutamine. To determine if physiological glucose and glutamine levels alter HCMV production, we measured infectious virus production at 120 hpi from MOI 1 or 3. Virus production was reduced by 2.8-logs at MOI 1 and 0.7-log at MOI 3 in DMEM with physiological levels of glutamine and glucose (**Fig 7F-G**).

**Fig 7.**
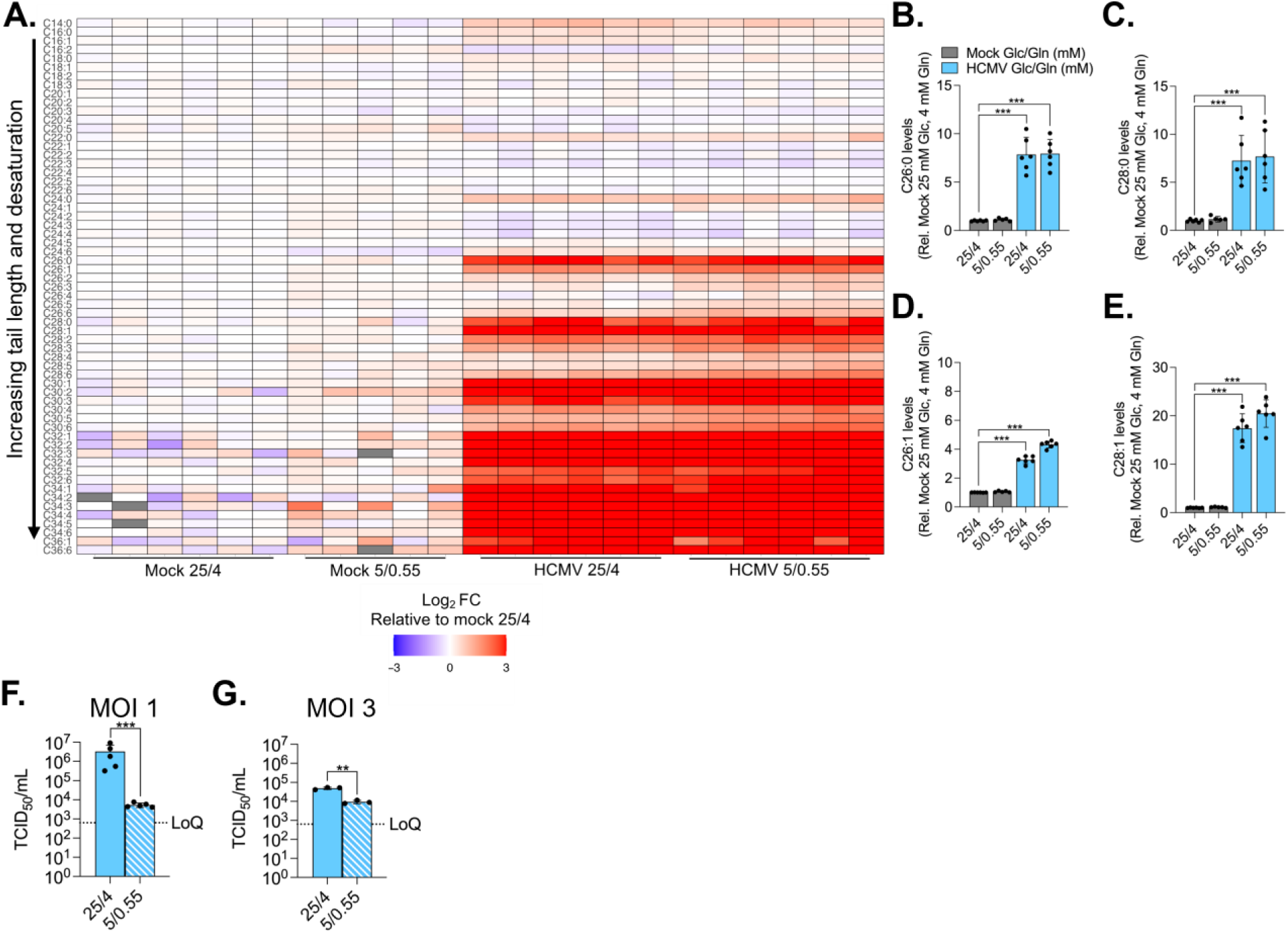
Physiological concentrations of glucose and glutamine support increased FA abundance during HCMV replication. (**A-E**) HFF-1 were infected with TB40/E-GFP or mock-infected. At 1 hpi, virus inoculum was removed and replaced with media containing 25 mM glucose and 4 mM glutamine (25/4) or 5 mM glucose and 0.55 mM glutamine (5/0.55). At 72 hpi, lipids were extracted and FA were saponified and measured using LC-MS. (**A**) Heatmap of FA abundance relative to mock 25/4. Each column is a replicate for the indicated condition. Grey bars indicate FA not identified in the sample. FA abundance for C26:0 (**B**), C28:0 (**C**), C26:1 (**D**), and C28:1 (**E**) relative to mock 25/4. (**F-G**) HFFs were infected with TB40/E-GFP or mock-infected at MOI 1 (**F**) or MOI 3 (**G**). Culture supernatant was collected at 120 hpi. Viral titers were measured by 50% tissue culture infection dose assay (TCID50). Error bars represent SD. Statistics were performed on transformed data for **F-G**. One-way ANOVA with Dunnett’s test were used to determine significance for **B-E**. Paired t-test was used to determine significance for **F-G**. *P* = 0.002, **, *P* < 0.001, ***. Error bars represent SD. *n*=3-6

Since physiological glucose and glutamine concentrations support HCMV-induced demand for FA synthesis, we evaluated the contribution of glutamine for FA synthesis in 5/0.55 DMEM. Specifically, we sought to understand if reducing glutamine and glucose would alter the use of glutamine in FA metabolism. We investigated this possibility with an isotopic labeling approach using U-^13^C-Gln in 5/0.55 DMEM compared to 25/4 DMEM. We infected or mock-infected HFFs and changed the culture media at 1 hpi to media containing either 4 mM or 0.55 mM U-^13^C-Gln and unlabeled glucose at 25mM or 5 mM, respectively. First, we examined if altering carbon source concentrations altered glutamine labeling of SFA and MUFA in mock-infected cells. We found the level of glutamine carbon use for FA in mock-infected cells was reduced in 5/0.55 compared to 25/4 for C14:0, C16:0, C20:1, and ≥C22 saturated and monounsaturated VLCFA (**Fig 8A-B**). For HCMV-infected cells, we also found reduced labeling in cells cultured in 5/0.55 compared to 25/4. Despite this reduction, we quantified a greater level of labeling in both DMEM conditions for HCMV-infected cells compared to mock-infected cells cultured in 25/4 for most SFA and ≥C26:1 MUFA. When directly comparing mock and infected cells cultured in 5/0.55, HCMV infection increased glutamine labeling of all SFA and ≥C24:1 MUFA compared to mock. The number of labeled carbons incorporated into C26:0 during infection was highest in 25/4 DMEM (**Fig 8C**). These observations indicate that the level of glutamine incorporation into FA depends on the concentration of glutamine and glucose. Moreover, the results demonstrate that glutamine flow to FA synthesis occurs independent of supraphysiological levels of glutamine and glucose.

**Fig 8.**
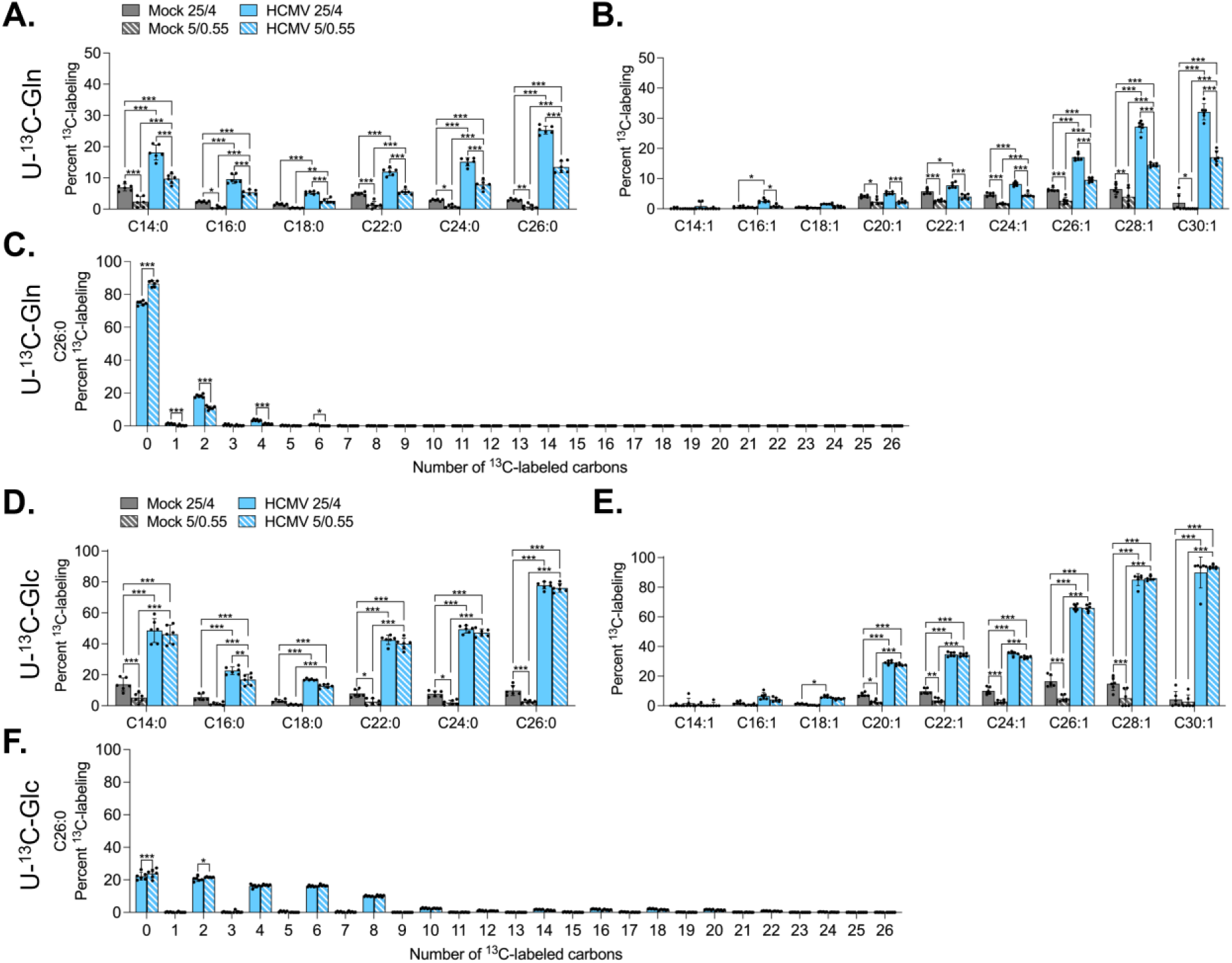
Glutamine and glucose contribute to FA synthesis when present at physiological levels during infection. HFFs were infected with TB40/E-GFP or mock-infected. At 1 hpi, virus inoculum was removed and replaced with media containing U-^13^C-Gln (**A-C**) or U-^13^C-Glc (**D-F**) at 25 mM glucose and 4 mM glutamine (25/4) or 5 mM glucose and 0.55 mM glutamine (5/0.55). At 72 hpi, lipids were extracted and FA were saponified and measured using LC-MS. Percent ^13^C-labeled SFA or MUFA were quantified for glutamine (**A-B**) or glucose (**D-E**) labeling. Percent ^13^C-labeled C26:0 in HCMV-infected cells for glutamine (**C**) or glucose labeling (**F**). Error bars represent SD. Two-way ANOVA with Šídák’s test were used to determine significance. *P* < 0.05, *; *P* < 0.01, **; *P* < 0.001, ***. *n*=6

Given our findings that reducing levels of glutamine and glucose in DMEM to human plasma-like concentrations resulted in a lower level of glutamine incorporation into FA, we asked if the flow of glucose to FA synthesis would be altered in 5/0.55 DMEM. For these experiments, we are using uniformly labeled glucose (U-^13^C-Glc) and unlabeled glutamine in 25/4 and 5/0.55 DMEM. We quantified the percent of FA containing at least one labeled carbon from glucose after correcting for natural isotopic abundance. We focused on glucose labeling of SFA and MUFA. In mock-infected cells, labeled C14:0, C20:1 and ≥C22 saturated and monounsaturated VLCFA were significantly reduced for 5/0.55 compared to 25/4 (**Fig 8D-E**). In contrast, HCMV-infected cells showed no difference in FA labeling when cultured in 5/.055 versus 25/4 except for a subtle but statistically significant reduction in C16:0 labeling. We counted the number of labeled carbons in C26:0 and found that ∼8-10 labeled carbons from glucose were incorporated at a similar level in HCMV-infected cells cultured in 25/4 or 5/0.55 (**Fig 8F**). These results indicate that the flow of glucose to FA synthesis at supraphysiological levels is maintained at physiological levels of glucose and glutamine during HCMV infection. Together, the metabolic tracing results using labeled glucose and glutamine in 5/0.55 DMEM suggest that glutamine and glucose carbon flow are, at least in part, differentially regulated since a difference was observed for glutamine but not glucose labeling.

## Discussion

Many viruses reprogram host metabolism to generate a metabolic environment favorable for viral replication. HCMV uses host factors to alter nutrient flow and remodel the lipidome for efficient virus production. Prior FA and lipidomic studies determined that HCMV reprograms glucose flow to supply carbon for lipogenesis (7, 10, 11, 27). Previous research demonstrated that HCMV replication is dependent on glutamine (16). Though it was hypothesized that glutamine could contribute to FA synthesis (5), the only demonstrated role of glutamine during HCMV replication was maintenance of the TCA cycle (6, 16). In this study, we determined that glutamine is metabolized via reductive carboxylation to feed FA and lipid synthesis; thereby, contributing to lipidome remodeling necessary for efficient virus production (**Fig 9**) (7, 10, 11). Moreover, our results suggest malic enzyme pathway may be active to support additional carbon flow to FA synthesis from both oxidative TCA and reductive carboxylation (**Figs 5, 6,** and **9**) (16). The contribution of glutamine to FA synthesis was likely overlooked until now since glucose is the primary carbon contributor to this process (**Fig 8**) (7). With this discovery, the field has now identified three carbon suppliers for FA synthesis during HCMV infection: glucose, glutamine, and acetate (6, 7, 10, 11). Why does the virus promote redundant carbon suppliers for the same process? HCMV has broad tissue distribution and cell tropism. The virus replicates in various nutrient niches in the body with vastly different metabolite availability than what is modeled in current cell culture approaches (40, 41). In the human body, it may be advantageous for HCMV to promote reprogramming of metabolic processes using different metabolite contributors to support the same molecular building blocks, e.g., FA and lipids. In this case, if metabolites are limited at the site of infection, another metabolite can compensate for loss. Our study highlights the importance of uncovering metabolic mechanisms that are overlooked in cell culture models but could be required for HCMV replication in the human body.

**Fig 9.**
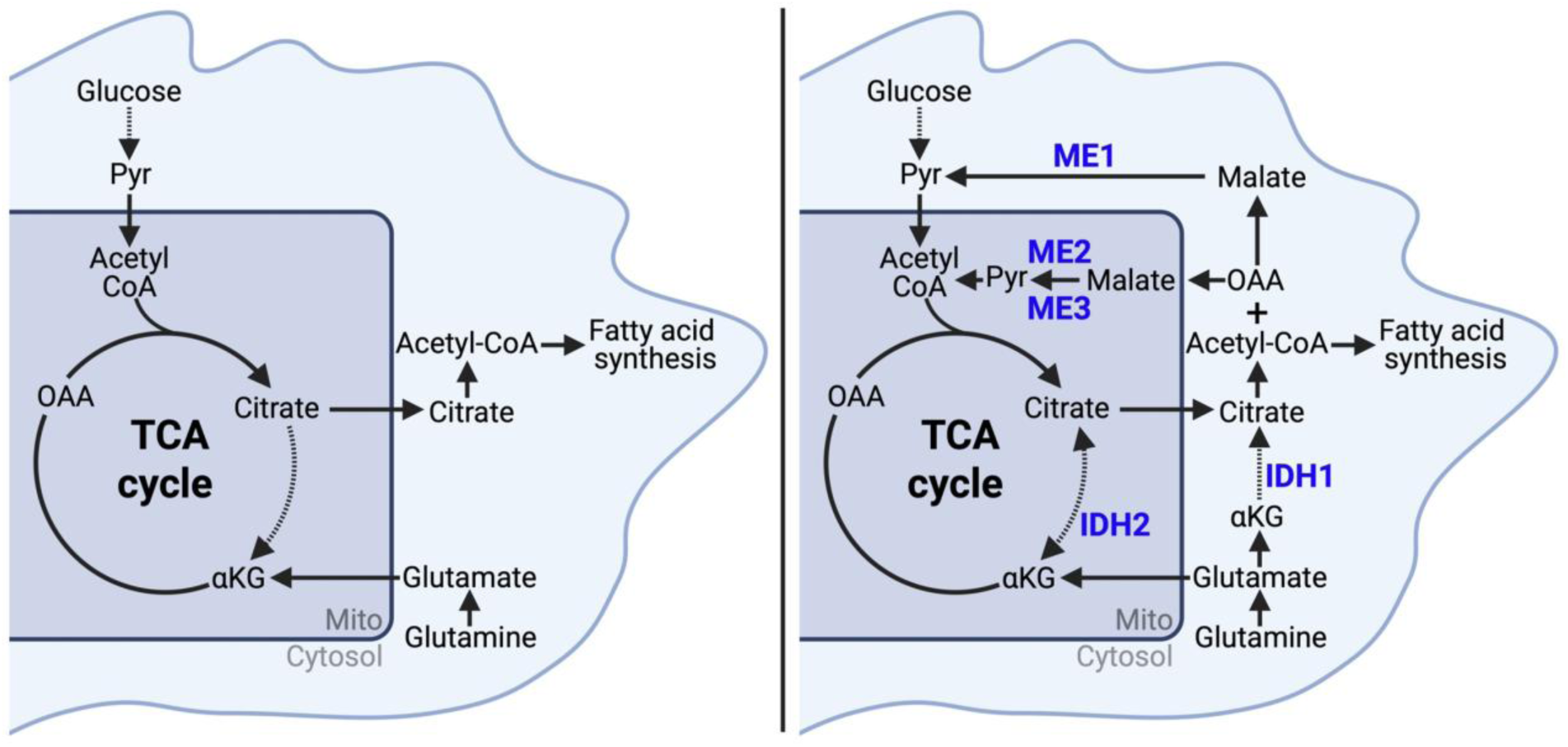
Model of glutamine carbon flow during HCMV replication. Pyruvate (Pyr); oxaloacetate (OAA); α-ketoglutarate (αKG). Dashed lines indicate metabolic intermediates not shown. (**Left**) Prior model of glutamine flow during infection. Glutamine is converted through glutaminolysis to αKG that feeds oxidative TCA cycle. (**Right**) Revised model of glutamine flow during infection. Glutamine is metabolized to glutamate and then to αKG in the cytosol. Alternatively, glutamate can enter the mitochondria and be metabolized to αKG. αKG is metabolized to isocitrate in the cytosol via IDH1 or the mitochondria via IDH2. Isocitrate is converted to citrate. Citrate produced through reductive carboxylation or oxidative TCA is cleaved via ATP citrate lyase to produce acetyl-CoA and OAA. OAA can be metabolized to malate and then converted to pyruvate via ME1 resulting in cytosolic pyruvate which can then be transported into the mitochondria for acetyl-CoA production. Alternatively, the malate to pyruvate reaction can occur in the mitochondria via ME2/3 resulting in mitochondrial pyruvate production and subsequent conversion to acetyl-CoA.

Our previous understanding of glutamine during HCMV replication was to maintain the TCA cycle through glutaminolysis, i.e., catabolism of glutamine to α-ketoglutarate that directly feeds the TCA cycle (6, 16). Our data demonstrate that glutamine is also metabolized through reductive carboxylation to support lipid synthesis in addition to its role in TCA cycle maintenance (**Figs 1**, **4, 9, S1,** and **S3**). These observations lead to a central question: Is there a metabolic decision to drive glutaminolysis versus reductive carboxylation, or are both roles of glutamine essential for HCMV replication (**Fig 9**)? This question cannot be easily answered with our current understanding. Glutamine deprivation starves HCMV-infected cells of both roles of glutamine. Chambers et al. demonstrated that α-ketoglutarate, oxaloacetate, and pyruvate supplementation are each sufficient to restore virus production during glutamine deprivation (16). Each supplement supports both metabolic pathways. α-Ketoglutarate can feed both reductive carboxylation and glutaminolysis, while oxaloacetate is a product of both pathways via cytosolic citrate cleavage (**Fig 9**). Pyruvate can be produced from glucose through glycolysis or from glutamine via the malic enzyme pathway and can also be converted into oxaloacetate via pyruvate carboxylase in specific metabolic conditions. It is possible the virus has evolved to promote glutaminolysis and reductive carboxylation to support both the TCA cycle and FA synthesis in specific metabolic niches where nutrients may be limiting (16). It remains to be investigated how these two processes are regulated and if alternative nutrients environments alter their regulation and contribution to metabolic needs during HCMV infection. Though our study and the Chambers et al. study cannot elucidate the essentiality of glutaminolysis versus reductive carboxylation, they highlight the remarkable adaptability of HCMV to different metabolic conditions.

Our data demonstrate that reductive carboxylation is the primary pathway through which glutamine contributes to FA synthesis (**Figs 2, 4, 5, 9, S2** and **S3**). Reductive carboxylation is typically active in hypoxic conditions or cancer cells (18–20, 31, 42). Increased reductive carboxylation activity has also been observed during adenovirus type 5 (Ad5) (non-enveloped, DNA virus) and murine norovirus (MNV) (non-enveloped, RNA virus) infection (43, 44). These studies identified an increase in 3-labeled TCA cycle metabolites during infection after culture in media containing uniformly labeled glutamine. Malate, fumarate, or aspartate with 3-labeled carbons are only obtained through reductive carboxylation of U-^13^C-Gln after citrate cleavage to acetyl-CoA and oxaloacetate. These studies did not directly measure glutamine carbon incorporation to FA, but it is plausible that acetyl-CoA production from reductive carboxylation could be used for FA synthesis during Ad5 and MNV replication. Reductive carboxylation and its contribution to the pro-viral metabolic environment is only beginning to be interrogated. Our results from HCMV, coupled with studies on Ad5 and MNV, indicate that reductive carboxylation is a shared metabolic mechanism in a diverse range of enveloped and non-enveloped DNA and RNA viruses.

Reductive carboxylation is active in cancer cells during hypoxia or impaired cellular respiration to sustain FA synthesis for rapid cellular proliferation (18–20, 31, 42). Conversion of α-ketoglutarate to isocitrate during reduction carboxylation is catalyzed by IDH1 (cytosolic) or IDH2 (mitochondrial) (**Fig 3A**). In cancer cells, both IDH1 and IDH2 are shown to support reductive carboxylation (18, 20). We found that IDH1 and IDH2 are expressed in HFFs but do not increase during HCMV infection (**Fig 3B-D**). These results suggest that reductive carboxylation during HCMV replication is regulated by other means, such as increased substrate availability (i.e., α-ketoglutarate) or IDH1 or IDH2 post-translational modification rather than increased enzyme expression (45, 46).

Our findings indicate HCMV-induced reductive carboxylation occurs through either cytosolic IDH1 or mitochondrial IDH2 activity (**Fig 9**). Though it is possible this activity can be active in both locations, our understanding of metabolic flux during HCMV replication suggests that reductive carboxylation occurs in the cytosol through IDH1 activity. Munger et al. demonstrated increased flux through oxidative TCA cycle during infection with glucose carbons primarily shuttled to citrate production for FA synthesis (6, 7, 10, 11). Chamber et al. showed glutamine supports oxidative turning of the TCA, which allows glucose shuttling to citrate production (6, 16). In support of these studies, we observed increased IDH3A levels (**Fig 3B** and **E**), suggesting that oxidative TCA cycle activity is supported by increased IDH3 that converts isocitrate to α-ketoglutarate. Together, these results indicate HCMV replication drives oxidative turning of the TCA cycle. This conclusion suggests that IDH2 in the mitochondria does not support the reverse reaction, i.e., α-ketoglutarate to isocitrate (**Fig 9**). If IDH1 is active in the cytosol, it would allow both mechanisms of glutamine use during infection: oxidative TCA cycle maintenance and reductive carboxylation.

Reductive carboxylation is the primary pathway through which glutamine contributes to FA synthesis (**Figs 2, 4, 5, S2** and **S3**). Comparison of U-^13^C-Gln and 5-^13^C-Gln FA labeling indicate that ∼30% percent of FA labeling comes from other metabolic means (**Fig 5C-F**). Cytosolic oxaloacetate that is produced from reductive carboxylation and oxidative TCA cycle can feed malic enzyme pathway, where it is converted to malate and pyruvate through malate dehydrogenase and malic enzyme, respectively (**Fig 6A-B**). The resulting pyruvate could feed oxidative TCA cycle to produce citrate to be used for FA synthesis. Malic enzyme pathway would support the increased FA labeling we observed between U-^13^C-Gln and 5-^13^C-Gln (**Fig 5C-D**). In support of this mechanism, ME activity is increased during HCMV infection (16), and we found all three isoforms of malic enzyme are increased during HCMV replication (**Fig 6C-F**). Malic enzyme activity could occur in the cytosol (ME1) or the mitochondria (ME2/ME3) (**Fig 9**). All three ME isoforms also produce NADPH, a cofactor for lipid synthesis, during malate to pyruvate conversion. HCMV could promote ME expression to support carbon flow to FA or to provide additional NADPH for lipid synthesis. Understanding the dynamic activity of enzymes that catalyze the same reaction in different cellular compartments is crucial for understanding metabolic reprogramming during infection. Our future studies will focus on elucidating the regulation and activity of ME isoforms and their role in HCMV replication.

HCMV infection remodels the lipid composition of the cell. We have previously demonstrated that several phospholipids and neutral lipid classes are increased during infection (7, 8). Our data indicates that glutamine carbons contribute to synthesis of all the lipid classes we examined except PG (**Fig 1A-B**). Moreover, glutamine was enriched in various tail lengths and degrees of saturation for PC, PE, DG, and TG (**Fig 1C-F**). Of the different phospholipid classes, PC with VLCFA tails (PC-VLCFA) are among the most increased by infection (7, 9). Consistent with these findings, PC show the greatest increase in labeling compared to mock by 72 hpi, and carbons from glutamine are enriched in VLCFA (**Figs 1A-B, 2B-C, S1A**). PC with saturated and monounsaturated fatty acyl tails showed the highest amount of labeling in HCMV-infected cultures compared to other phospholipid classes (**Fig 1C-F**, and **S1**). These data indicate that glutamine supports SFA and MUFA production to support increased PC-VLCFA synthesis (9–11). Overall, our results indicate glutamine contributes to lipidome remodeling by supporting FA and lipid synthesis.

HCMV infection promotes reductive carboxylation to supply glutamine carbons for FA and lipid synthesis (**Figs 1, 2, 4, S1, S2,** and **S3**), indicating a viral protein drives glutamine metabolic reprogramming. HCMV UL38 is heavily involved in metabolic reprogramming during HCMV infection (11, 23, 39). UL38 supports fatty acid elongation and drives consumption of glucose and glutamine (11, 39). Moreover, UL38 is necessary and sufficient for glutamine consumption (39). Raymonda et al. demonstrated that cells expressing UL38 increased glutamine consumption in response to glucose deprivation (23). Notably, acetyl-CoA and malonyl-CoA, carbon suppliers for FA synthesis, were increased in UL38 expressing cells deprived of glucose (23). It is plausible UL38 could drive glutamine carbon flow to FA synthesis via reductive carboxylation. UL37×1 is another potential candidate that may induce glutamine metabolic reprogramming since we have demonstrated UL37×1 supports lipidome remodeling during infection (7). While UL37×1 is not sufficient to induce lipid synthesis on its own (7), it could function with UL38 to regulate metabolism, including glutamine feeding reductive carboxylation for FA and lipid synthesis.

Collectively, our study adds additional information to our current model of HCMV metabolic flow during infection (**Fig 9**). First, HCMV infection promotes glutamine carbon flow through reductive carboxylation to FA synthesis, and second, infection increases the activity in metabolic steps that feed additional glutamine carbons to FA production, likely through increased malic enzyme levels. Moreover, these results suggest that HCMV-induces metabolic reprogramming to support FA synthesis from several metabolic pools, allowing metabolic flexibility in different nutrient niches at sites of infection. Overall, our results expand understanding of the role of glutamine and metabolic reprogramming during viral infection.

## Materials and Methods

### Cells, viruses, and reagents

Human foreskin fibroblasts (HFFs) or HFF-1 (American Type Culture Collection; ATCC) cells were cultured in Dulbecco’s modified Eagle’s medium (DMEM) containing 4.5 g/L (25 mM) glucose (Gibco 11965), 10% fetal bovine serum (FBS), 10 mM HEPES, and penicillin/streptomycin (P/S). Prior to infection, cells were held at confluency for three days in full growth medium and then another day in serum-free (SF) medium (DMEM, 10 mM HEPES, P/S). DMEM containing ^13^C-labeled glucose or glutamine were prepared using DMEM powder (Gibco) and adding 44 mM sodium bicarbonate with the indicated concentration of ^13^C-labeled glucose or glutamine and the indicated concentration of unlabeled glucose or glutamine.

Cells were infected with HCMV strain TB40/E encoding free GFP (TB40/E-GFP) at MOI 3 IU/cell (unless noted otherwise) for 1 h and washed two times with PBS before adding fresh media. Mock-infected cells were treated with inoculum that did not contain virus. At 48 hpi, media was changed to maintain nutrient levels.

Virus stocks were produced by propagating BAC-derived HCMV in fibroblasts and concentrating infected cell culture supernatant by pelleting through a sorbitol (20% sorbitol, 50 mM Tris pH 7.2, 1 mM MgCl2) cushion for 80 min at 52,931xg using ultracentrifugation. Viruses were resuspended in SF DMEM. Stocks and experimental samples were titered using 50% tissue culture infectious dose assay (TCID50) beginning with a 1:10 dilution of concentrated virus. GFP-positive wells were counted at two-weeks post infection on duplicate plates. A well was considered positive if it contained a foci of ≥3 GFP-positive cells.

### Lipid Analysis

Lipid analysis was performed using LC-MS/MS. Duplicate wells for each condition were analyzed, and a third well was used to determine a cell count for normalization. Additional wells that did not contain cells were used as a control to determine contaminants from the extraction or LC-MS/MS steps. Cells were washed three times in cold PBS and fixed in cold 50% methanol (MeOH) with 0.1 M HCl. Samples were scraped and transferred to glass vials. Lipids were extracted two times in chloroform and dried under nitrogen gas. Lipids were resuspended in 1:1:1 solution of Methanol:Isopropanol:Chloroform at 100 µl per 200,000 cells, stored at 7°C in an autosampler, and separated using reverse-phase chromatography on a Vanquish UHPLC system using a Kinetex C18 reverse-phase column (Phenomenex) at 60°C using solvent A (40:60 water:MeOH, 10 mM ammonium formate, 0.1% formic acid) and solvent B (10:90 MeOH:Isopropanol, 10 mM ammonium formate, 0.1% formic acid). UHPLC was performed at a flow rate of 0.25 mL/min with a total run time of 30 min. The gradient was established at 25% solvent B for 2 min, 65% solvent B at curve value 4 from 2-4 min, 65% held for 4-5 min, 100% solvent B at a curve of 4 from 5-16 min, and held at 100% solvent B from 16-20 min. The column was briefly washed and equilibrated after each sample. Lipids were analyzed using a Thermo Fisher Q-Exactive Plus mass spectrometer operating in full MS1/dd-MS2 with TopN mode set to 5 scans. MS1 spectra were collected from 200 to 1,600 *m/z* at 70,000 resolution using an automatic gain control (AGC) target of 1e6 with transient time of 250 ms. Lipids were ionized using HESI in positive and negative mode. In positive mode, 8 µl of sample was analyzed using sheath gas of 45, auxiliary gas of 16, sweep gas of 2, spray voltage of 3.5 kV, S-lens RF of 65, ion transfer tube temperature of 320°C, and vaporization temperature of 220°C. In negative mode, 10 µl of sample was analyzed using sheath gas of 44, auxiliary gas of 14, sweep gas of 2, spray voltage of 3.3 kV, S-lens RF of 76, ion transfer tube temperature of 320°C, and vaporization temperature of 220°C. MS2 spectra were collected at 35,000 resolution with an AGC target of 1e5. In positive mode, NCE of 30 was used. In negative mode, NCE value of 20 was used. The instrument was calibrated weekly and immediately prior to analysis. For all lipid analysis, blank samples were run before, after, and interspersed between samples. Data were analyzed using Maven 2 software (47) and corrected for the natural ^13^C abundance using MATLAB (6). The percent of labeled carbons were determined by subtracting the unlabeled fraction from one and multiplying by 100 to obtain percent ^13^C label.

### Fatty acid analysis

Fatty acid analysis was performed using LC-MS. Duplicate wells for each condition were extracted and analyzed in parallel. Additional wells that did not contain cells were used as a control to determine contaminants from the extraction or LC-MS steps. Cells were collected and lipids extracted as detailed in the lipid analysis section. After drying under nitrogen gas, FA tails were chemically cleaved by saponifying in 90% MeOH and 0.3 M potassium hydroxide at 80°C for 1 h followed by neutralization using formic acid. FA were extracted two times in hexane and dried under nitrogen gas. FA were resuspended in 600 µl of 1:1:1 solution of MeOH:Isopropanol:Chloroform and stored at 7°C in an autosampler. FA were separated by reverse-phase chromatography on a Vanquish UHPLC system using a Luna C8 reverse-phase column (Phenomenex) at 60°C and solvent A and solvent B described in the lipids analysis section. UHPLC was performed at a flow rate of 0.2 mL/min with a total run time of 30 min. The gradient was established at 10% solvent B for 2.5 min, 100% solvent B at curve value 5 from 2.5-10 min, and held at 100% solvent B from 10-22 min. The column was briefly washed and equilibrated after each sample. FA were analyzed using a Thermo Scientific Orbitrap Exploris 240 mass spectrometer at a resolution of 120,000 for unlabeled FA analysis or 240,000 for ^13^C-labeled FA analysis. Spectra were collected at a range of 200-650 *m/z*. FAs were ionized by HESI in negative mode using 10 µl of sample. The following settings were used: sheath gas of 50, auxiliary gas of 25, sweep gas of 10, spray voltage of 3 kV, RF lens of 110%, ion transfer tube temperature of 300°C, and vaporization temperature of 320°C. The internal calibration source “Easy-IC” was used in Scan-to-Scan mode. For labeling experiments, the instrument was calibrated immediately prior to the start of the analysis; otherwise, the instrument was calibrated weekly. Blank samples were run before, after, and interspersed between samples. Data were analyzed using Maven 2 software (47). For labeling experiments, the data were corrected for the natural ^13^C abundance using MATLAB (6). Unlabeled FA abundances were determined by calculating the total ion count for each sample and setting each FA in the sample relative to the total ion count. The percent of labeled carbons were determined by subtracting the unlabeled fraction from one and multiplying by 100 to obtain percent ^13^C label.

### Protein Analysis

Protein levels were analyzed by western blot. Whole-cell lysates were collected by in-well lysis followed by scraping. Proteins were resolved by SDS-PAGE using Mini-Protean 4-20% gradient gels (BioRad) and transferred to nitrocellulose membrane (LI-COR). Membranes were blocked in 5% milk in tris buffered saline with 0.1% Tween 20 (TBST). Primary antibodies were diluted in 5% milk in TBST and incubated with rocking for 1 h at room temperature or overnight at 4°C. Secondary antibodies were diluted in 5% milk in TBST and incubated for 1 h with rocking in the dark at room temperature. Images were taken on a LI-COR Odessey CLx imager, and quantitation was performed using Image Studio software.

The following primary antibodies were used for western blot analysis: Mouse-anti tubulin (Sigma; 1:1,000), mouse anti-IE1 (clone 1B12; 1:100) (11), rabbit anti-IDH1 (proteintech; 1:1,000), rabbit anti-IDH2 (proteintech; 1:1,000), rabbit anti-IDH3A (proteintech;1:1,000), rabbit anti-ME1 (proteintech; 1:1,000), rabbit anti-ME2 (proteintech; 1:1,000), and rabbit anti-ME3 (ThermoFisher; 1:1,000). HCMV IE1 antibody was a gift from Dr. Thomas Shenk (Princeton University). Goat anti-mouse DyLight 800 (1:10,000) or goat anti-rabbit DyLight 680 (1:10,000) were used for secondary antibodies.

### Graphics and Statistics

Schematics and models were created with BioRender.com. Graphs were designed and statistical testing were performed using GraphPad Prism. The appropriate statistical tests for each experiment are noted in the figure legends. Heatmaps were designed using R.

## Acknowledgments

We thank Dr. Jim Alwine for helpful feedback. We thank members of the Purdy, Goodrum, and Van Doorslaer laboratories for their input on the project. Research reported in this publication was supported by the National Institute of Allergy and Infectious Diseases of the National Institutes of Health under Award Number R01AI162671 (J.G.P.), R01AI155539 (J.G.P.), and F32AI178919 (R.L.M.). The content is solely the responsibility of the authors and does not necessarily represent the official views of the National Institutes of Health. Additional funding was provided by the BIO5 Institute Postdoctoral Fellowship program awarded to R.L.M.

---

**Fig S1.**
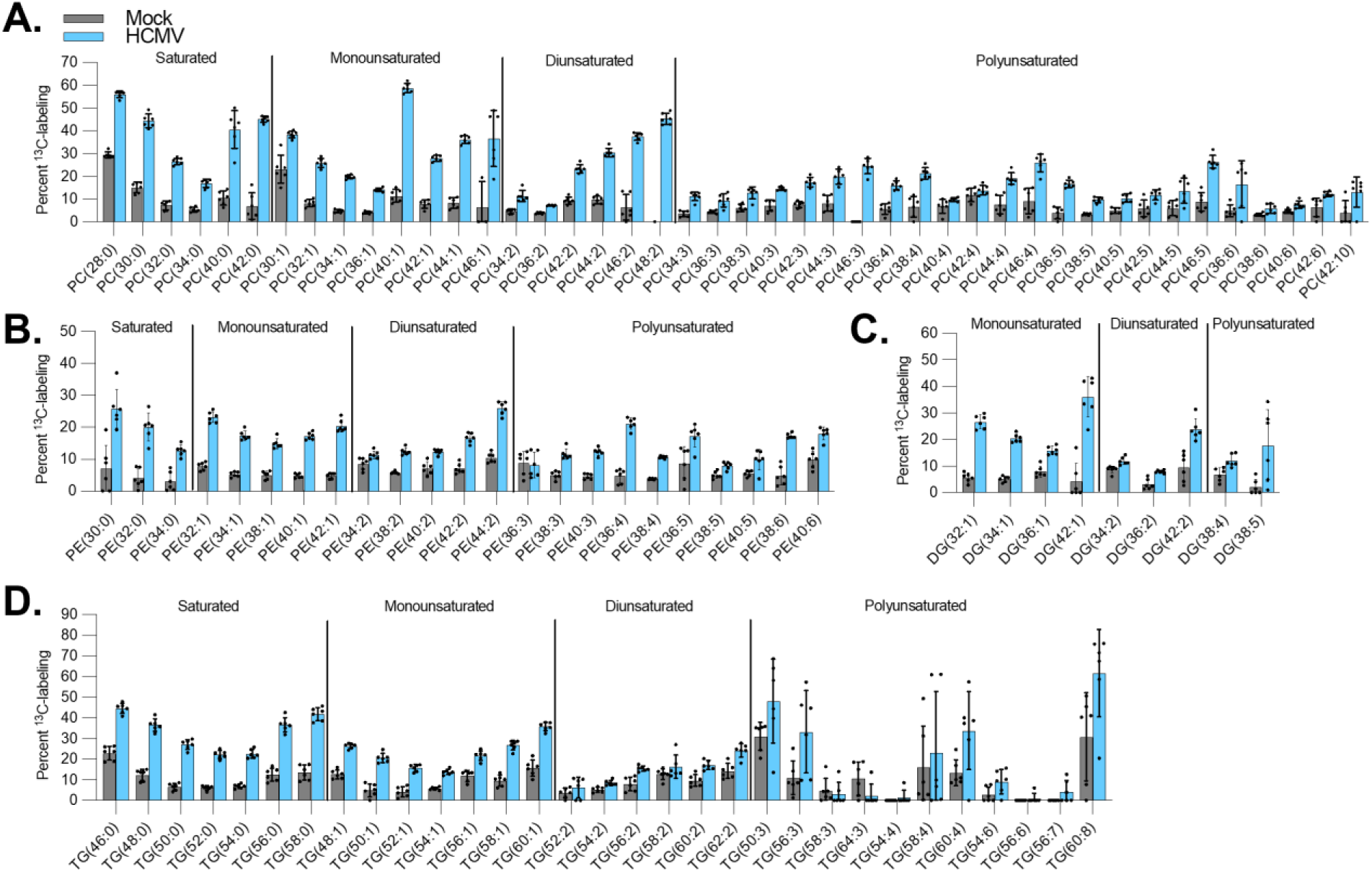
Glutamine supports lipid synthesis across tail desaturations during HCMV replication. HFFs were infected with TB40/E-GFP or mock-infected. At 1 hpi, virus inoculum was removed and replaced with media containing uniformly labeled ^13^C glutamine (U-^13^C-Gln). At 72 hpi, lipids were extracted and measured using LC-MS/MS. Percent labeled lipids are shown for PC (**A**), PE (**B**), DG (**C**), and TG (**D**). Error bars represent SD. *n*=6

**Fig S2.**
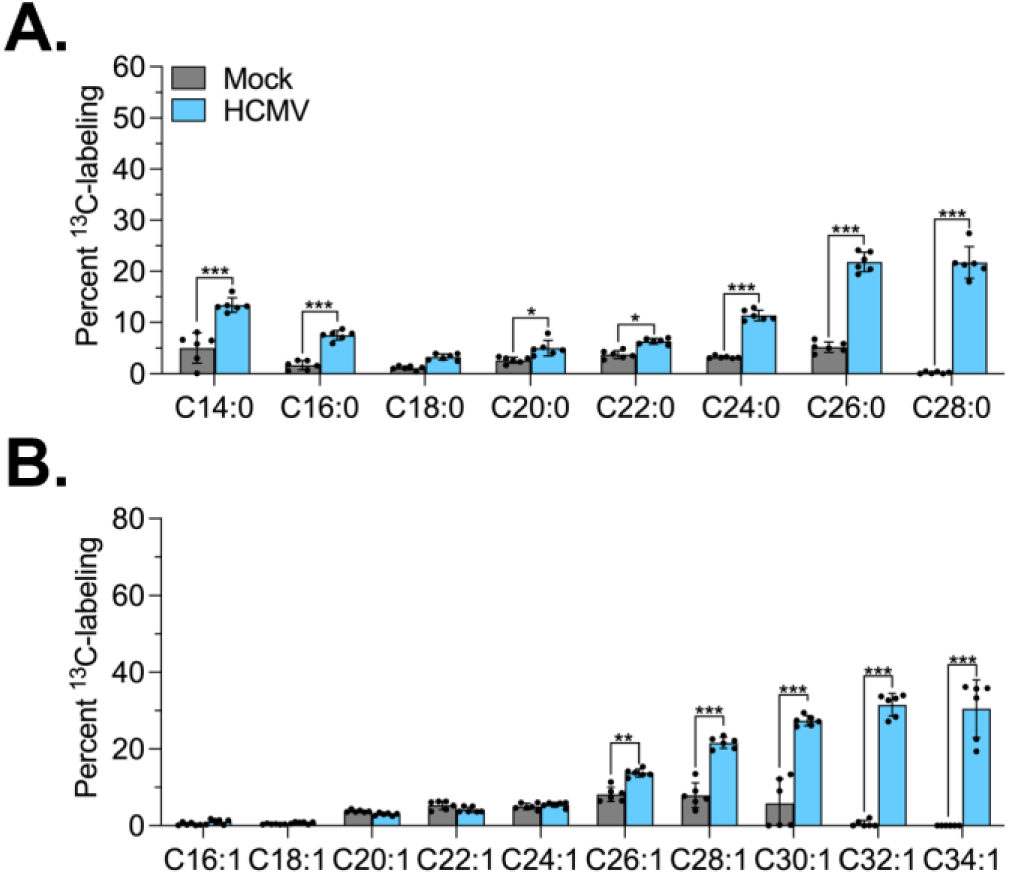
Glutamine supports FA synthesis by 48 hpi during HCMV replication. (**A-C**) HFFs were infected with TB40/E-GFP or mock-infected. At 1 hpi, virus inoculum was removed and replaced with media containing U-^13^C-Gln. At 48 hpi, lipids were extracted and FA were saponified. Labeled FA were measured using LC-MS. Percent ^13^C-labeled SFA (**A**) and MUFA (**B**) were quantified. Error bars represent SD. Two-way ANOVA with Šídák’s test were used to determine significance. *P* < 0.05, *; *P* < 0.01, **; *P* < 0.001, ***. *n*=6

**Fig S3.**
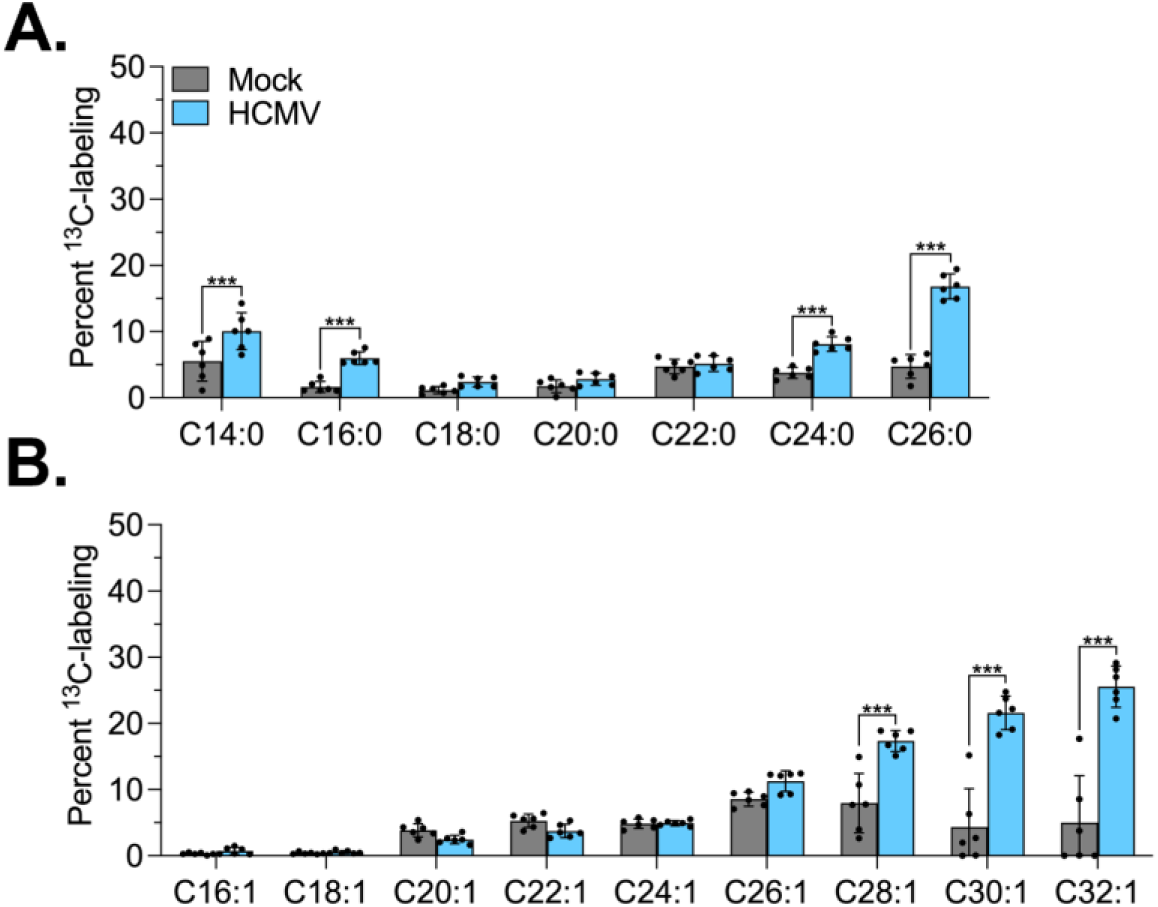
Reductive carboxylation activity increases by 48 hpi during HCMV replication. (**A-C**) HFFs were infected with TB40/E-GFP or mock-infected. At 1 hpi, virus inoculum was removed and replaced with media containing 5-^13^C-Gln. At 48 hpi, lipids were extracted and FA were saponified. Labeled FA were measured using LC-MS. Percent ^13^C-labeled SFA (**A**) and MUFA (**B**) were quantified. Error bars represent SD. Two-way ANOVA with Šídák’s test were used to determine significance. *P* < 0.001, ***. *n*=5-6

